# Quantifiable Blood TCR Repertoire Components Associated with Immune Aging

**DOI:** 10.1101/2024.07.19.604275

**Authors:** Jing Hu, Mingyao Pan, Brett Reid, Shelley Tworoger, Bo Li

## Abstract

T cell senescence results in decayed adaptive immune protection in older individuals, with decreased or increased abundance of certain T cell phenotypic subpopulations. However, no study has linked aging to the dynamic changes of T cell clones. Through a newly develop computational framework, Repertoire Functional Units (RFU), we investigated over 6,500 TCR repertoire sequencing samples from multiple human cohorts. Our analysis identified age-associated RFUs repeatedly and consistently across different cohorts. Quantification of RFU decreases with aging revealed accelerated loss under immunosuppressive conditions. Systematic analysis of age-associated RFUs in clinical samples manifested a potential link between these RFUs and improved clinical outcomes during acute viral infections, such as lower ICU admission and reduced risk of developing complications. Finally, our investigation of bone-marrow transplantation patients indicated a secondary expansion of the age-associated clones upon receiving stem cells from younger donors. Together, our results suggest the existence of certain clones or a ‘TCR clock’ that could reflect the immune functions in aging populations.

## Introduction

Deterioration of the immune system is one of the key features of aging^1^. Previous work has uncovered systematic changes in the innate and adaptive cellular compartments in older individuals^2,3^. Many studies have further focused on T cell senescence given their critical role in host defense against external pathogens and cancer^4,5^. Phenotypically, exhaustion-like T cells emerge and accumulate in aged tissues, with an altered, pro-inflammatory cytokine production program^6^. Quantitatively, there is continuous shrinkage of naïve T cell populations due to thymic atrophy^7,8^, which is usually accompanied by the expansion of memory clones in older individuals^9^. In addition, decreased abundance of innate-like T cell compartments was also observed, such as *γδ* T cells^10^ and mucosal-associated invariant T cells^11^. Consistently, studies using the immunogenomics profiling of T cell receptors (TCR) confirmed a robust decrease in repertoire diversity with aging^9,12–14^. Concomitant with the reduction of repertoire diversity, certain clones become more dominant while others decline or disappear, resulting in imbalanced naïve-memory ratio and increased clonality^13,15,16^.

Compared to cross-sectional studies, research based on longitudinal cohorts could estimate age-associated changes more precisely given the inter-individual heterogeneity of TCR repertoires^17^. Yoshida et al. profiled TCRβ repertoires of CD4+ and CD8+ T cells collected from six healthy donors over three visits, and reported significantly decreased CDR3β diversity and increased frequencies of clonal populations with age for CD8+ T cells^18^. Another longitudinal cohort with more donors and larger age spans further confirmed the reduction of richness in the naïve CD4+ and CD8+ subsets, but not in memory T cells^19^. Contrarily, retention of the same TCRs is found in different timepoints of the same donors, with more prominence in CD8+ memory cells than in CD4+ or naïve cells^19^.

While these studies provided critical insights on the overall decay of adaptive immune functions in aging populations, there has been no report of the age impact on specific T cell clones defined by the T cell receptor sequences. Such an investigation is challenging given the vast diversity of the human TCR repertoire^20^, yet it holds the promise to uncover shared ‘aging antigens’ that could be targeted in the older population for improved immune protection^6^. In this work, we seek to address this challenge with our newly developed TCR analysis method, Repertoire Functional Unit (RFU)^21^, which dissects the hypervariable TCR repertoire into quantifiable segments, or RFUs, that can be compared across different individuals. We applied this approach to investigate a large number of repertoire samples deeply sequenced for the β chain complementarity determining region 3 (CDR3) of TCRs and systematically studied the age associations of individual RFUs. Our analysis revealed 13 RFUs that showed dynamic changes with age and confirmed this observation in multiple independent cohorts. Through investigation of the clinical impact of these RFUs, we confirmed their prognostic value during acute viral infections and quantified their loss under normal or pathological conditions. Our work provided novel insights over immune aging that can open up potential opportunities for its reversal.

## Results

### Robustly age-associated TCR signatures in multiple human cohorts

We first implemented Repertoire Functional Unit, or RFU, our recent computational method to analyze the TCR repertoire samples^21^. In brief, each TCR sequence was projected to a 500-dimensional Euclidean space, with shorter distance between a pair of TCRs representing higher similarity. The embedding space was then divided into 5,000 conservative TCR neighborhoods, with the centroid of each neighborhood defined as an RFU. A new TCR-seq sample can be converted into a 5,000-length numeric vector of RFU counts, which can be compared across multiple samples in a large cohort.

We then investigated the age associations of RFUs using three TCR-seq blood sample cohorts, including Nolan cohort^22^ of 1,414 adults with COVID-19, Emerson cohort of 666 samples mainly from healthy adults^23^ and Mitchell cohort of 359 samples from children^24^. RFU-by-sample matrix (hereby denoted as RFU matrix) of each cohort was calculated using the above method. We compared the age associations of the two adult cohorts and observed consistency of a subset of negatively correlated RFUs in both cohorts (**Figure 1a, Table S1**). We defined the 13 RFUs with Spearman’s ρ≤ −0.2 in the Emerson cohort and ρ≤ −0.3 in the Nolan cohort as ‘essential RFU’, or eRFU (**Figure 1b** and **Figure S1a**). The T cell clones of these eRFUs demonstrated an approximately 50% reduction by age 80 or above compared to early adulthood (**Figure 1b**). This trend was reversed when we investigated the Mitchell cohort of younger individuals: all the adult age-related eRFUs were positively associated with age between birth to early 20s (**Figure 1c-d**). Combining adult and children cohorts, we obtained the full spectrum of non-linear age association (**Figure 1e**). All the 13 eRFUs exhibited dynamic changes with chronological age, experiencing rapid expansion after birth. They reached peaks around age 27 and then gradually decreased, ultimately returning to the birth level as participants aged.

**Figure 1.**
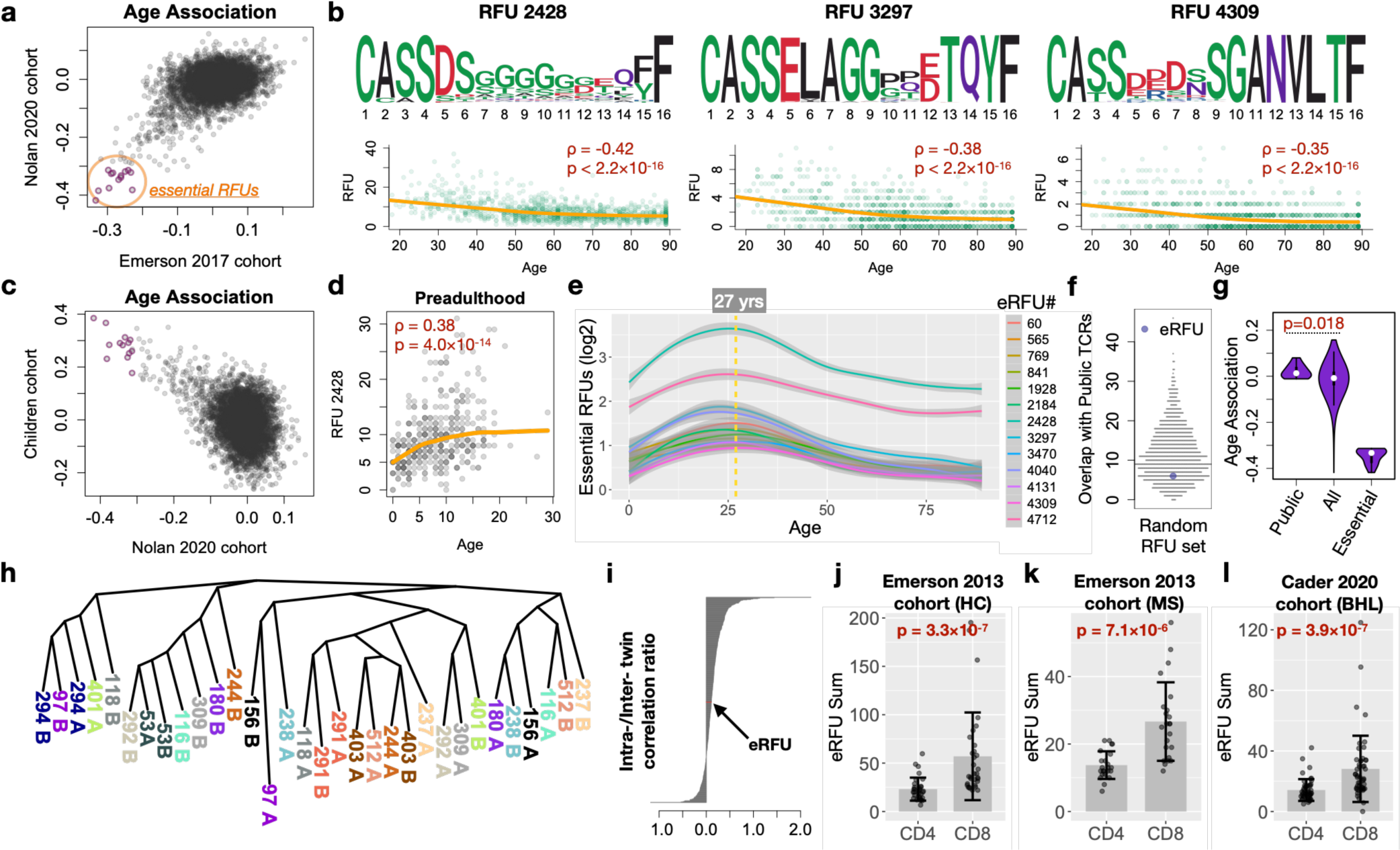
Identification and characterization of age-associated essential RFUs (eRFUs). **a**) Scatter plot showing the aligned age associations calculated from two large sample cohorts. Selection criteria of eRFUs were described in the main text. RFUs were defined by mapping the top 10,000 most abundant TCR clones in a repertoire to the embedding space. Statistical significance was evaluated using Spearman’s correlation test, with FDR corrected using the Benjamini-Hochberg approach. **b**) Sequence logo plot paired with scatter plot showing the direct age association for 3 eRFUs. **c**) Scatter plot showing the anti-aligned age associations from one adult and one childhood cohort, with eRFUs labeled with red circles. **d**) Positive age association of a selected eRFU in the childhood cohort. Statistical significance of the association in b) and d) was evaluated using Spearman’s correlation test. **e**) Smoothed lines showing dynamic changes with age for all 13 eRFUs. Confidence intervals of the smooth curves were calculated using 95% standard error estimated by Loess polynomial fitting. **f**) Beeswarm plot showing the distribution of overlaps of public TCRs with each of 1,000 random RFU set. eRFU was colored in purple. **g**) Violin plots comparing age associations for 3 groups of RFUs. Statistical significance was estimated with one-way ANOVA. **h**) Neighbor joining tree showing the relationships between pairs of twins with distance matrix calculated using the 13 eRFUs. **i**) Barplot showing the ratio of intra-/inter-twin associations of random RFU sets or eRFU. Associations were calculated as Spearman’s correlation using RFU as markers between a pair of individuals. **j**-**l**) Boxplots showing enrichment of eRFUs into CD4 or CD8 subsets in 3 independent adult cohorts. Statistical significance was evaluated using two-sided Wilcoxon rank sum test.

We next explored the nature of eRFU T cells. Public TCRs, defined as TCRs shared across multiple individuals, appear to be an important part of the TCR repertoire^25^. To explore the relationship between eRFU and public TCRs, we repeated the previous work^12^ to define public TCRs. We found that eRFUs were not enriched for public TCRs compared to a randomly selected RFU set (**Figure 1f**) and the public TCRs were not significantly associated with age (**Figure 1g**). Next, to understand how eRFUs are shaped by genetic factors, we took advantage of a TCR-seq sample cohort of identical twins^26^. Interestingly, while the TCR repertoire is overall more similar within than between twins (**Figure S1b**), this heritability was not observed for the eRFUs (**Figure 1h**). Further, intra-twin over inter-twin similarity calculated as Spearman’s ρ between a pair of individuals using eRFUs were not higher when compared to those using randomly selected RFU sets (**Figure 1i**). Finally, we sought to determine whether eRFUs had a specific T cell phenotypic subset. We analyzed two sample cohorts with memory CD4 and CD8 T cell repertoires separately sequenced^27,28^ and discovered that eRFUs were consistently enriched for the CD8+ population (**Figure 1j-l**), suggesting that eRFUs clones are potentially cytotoxic. However, examination of their associations to the common MHC-I alleles did not identify any significant associations (**Figure S2a**). These observations suggested that eRFUs are non-public and MHC-independent CD8 T cells.

### MAIT signatures of T cells with eRFU receptors

Previous studies indicated that mucosal associated invariant T (MAIT) cells possess the TCR motifs observed in the eRFUs with Aspartic or Glutamic acid on the 5^th^ position^29^. Consistent with our observation, MAIT cells were not MHC-I restricted as they recognize antigens bound by MR1^30^. Interestingly, we identified one putative MAIT TCR, M33.64 that carries the sequence signatures of eRFU 841 (Figure S2b). M33.64 binds to 5-(2-oxopropylideneamino)-6-d-ribitylaminouracil, or 5-OP-RU (Figure S2c), a potent MAIT activator derived from vitamin B metabolism in bacteria^31^. Further, although MAIT cells are known for their invariant ɑ chain^32^, conserved patterns in the β chain have also been observed^29^, consistent with our findings. We therefore investigated a single cell RNA-seq dataset paired with TCR sequences from human T cells of diverse phenotypes^33^. T cells assigned to eRFUs were enriched in cluster 7 (**Figure S2d**), which overexpressed putative markers of MAIT cells (**Figure S2e**). We further selected all the 2,619 annotated MAIT cell TCRs and analyzed each of the eRFUs. 7 out of the top 40 RFUs enriched in the MAIT TCRs (O/E≥ 50) were eRFUs (**Figure S2f**). These results indicated that most eRFU T cells belong to a subset of CD8+ MAIT cells.

To further investigate the nature of eRFUs, we analyzed another recent scRNA-seq dataset^34^, which contained paired scTCR-seq samples from flow-sorted MAIT cells and conventional memory T cells (Tmem). We confirmed that eRFUs were dominantly expressed by MAIT cells (**Figure S3a**), with an enrichment in the TRAV1-2+ subset (**Figure S3b**). Unbiased differential gene expression analysis revealed more upregulated genes in the eRFU cells compared to the non-eRFU MAIT cells (**Figure S3c**), suggesting a globally different transcriptional program between the two groups. Particularly, putative T cell stemness markers TCF7, BACH2 and KLF2^35–37^ were significantly downregulated in the eRFU group (**Figure S3d**). Unbiased analysis using signature gene sets (up- or down-regulated in CD8 stem T cells) ^38^ confirmed a low stemness state of eRFU MAITs (**Figure S3e**). Finally, pseudotime analysis^39^ with direction determined by CytoTRACE^40^ revealed that eRFU cells are significantly enriched at the most-differentiated end of the trajectory (**Figure S3f-g**). These results collectively indicated that eRFU-expressing cells belong to a more differentiated group of CD8+ MAIT cells.

The same dataset contained individuals aged from 27 to 65, allowing for the investigation of how stemness of eRFU MAITs trends with age. First, T cell stemness markers (TCF7, BACH2, KLF2) ^35–37^ consistently decrease with age (**Figure S4a**). The activity of transcription factors BACH2 and KLF2, measured by the expression levels of their regulating targets by SCENIC^41^, showed a similar trend (**Figure S4b**). Signature genes upregulated CD8 stem cells decrease with age in eRFU MAITs, while an opposite trend was observed for the down-regulated genes (**Figure S4c-d**). These results potentially indicated that the faster shrinkage of eRFU clones than the other MAIT cells is likely due to their relatively lower and continuously decaying stemness with aging.

### Quantification of eRFU loss with age in adults

We next sought to quantify the rate of eRFU decrease in adult cohorts. First, we observed a linear, homoscedastic relationship between age and log-scaled total eRFU summation in adults (**Figure S5a**). In the Nolan cohort, log_10_ eRFU sum is estimated to decay at the rate of 0.0092 ± 0.00044 (mean ± s.d., **Figure 2a**), which is approximately 2.1% loss per year. The second discovery set, Emerson 2017 cohort, provided a similar estimation of −0.0088, *i.e.* 2.0% annual loss (**Figure 2b**). To confirm these results, we analyzed two additional adult patient cohorts: Greenberger 2022 cohort^42^ with 480 subjects and Britanova 2016 cohort with 32 individuals^12^. The Britanova cohort was profiled using a different data generation method based on RNA, while all the other cohorts were profiled using genomic DNA. Interestingly, we observed similar rate of decay in both cohorts, with −0.0076 (1.7%) and −0.011 (2.5%) respectively (**Figure 2c-d**). We observed similar trend using another large human adult cohort with categorical age information^43^, which saw a rate of −0.093 per 10 years on the log_10_ scale (**Figure 2e**). Further analysis of racial groups revealed a potentially faster eRFU loss among Asian and Pacific Islanders (**Figure S5b**), though statistical significance was not reached due to small sample size. These results suggested that total count of eRFUs decreases by approximately 2% each year in the general population aged from 30s till 80s.

**Figure 2.**
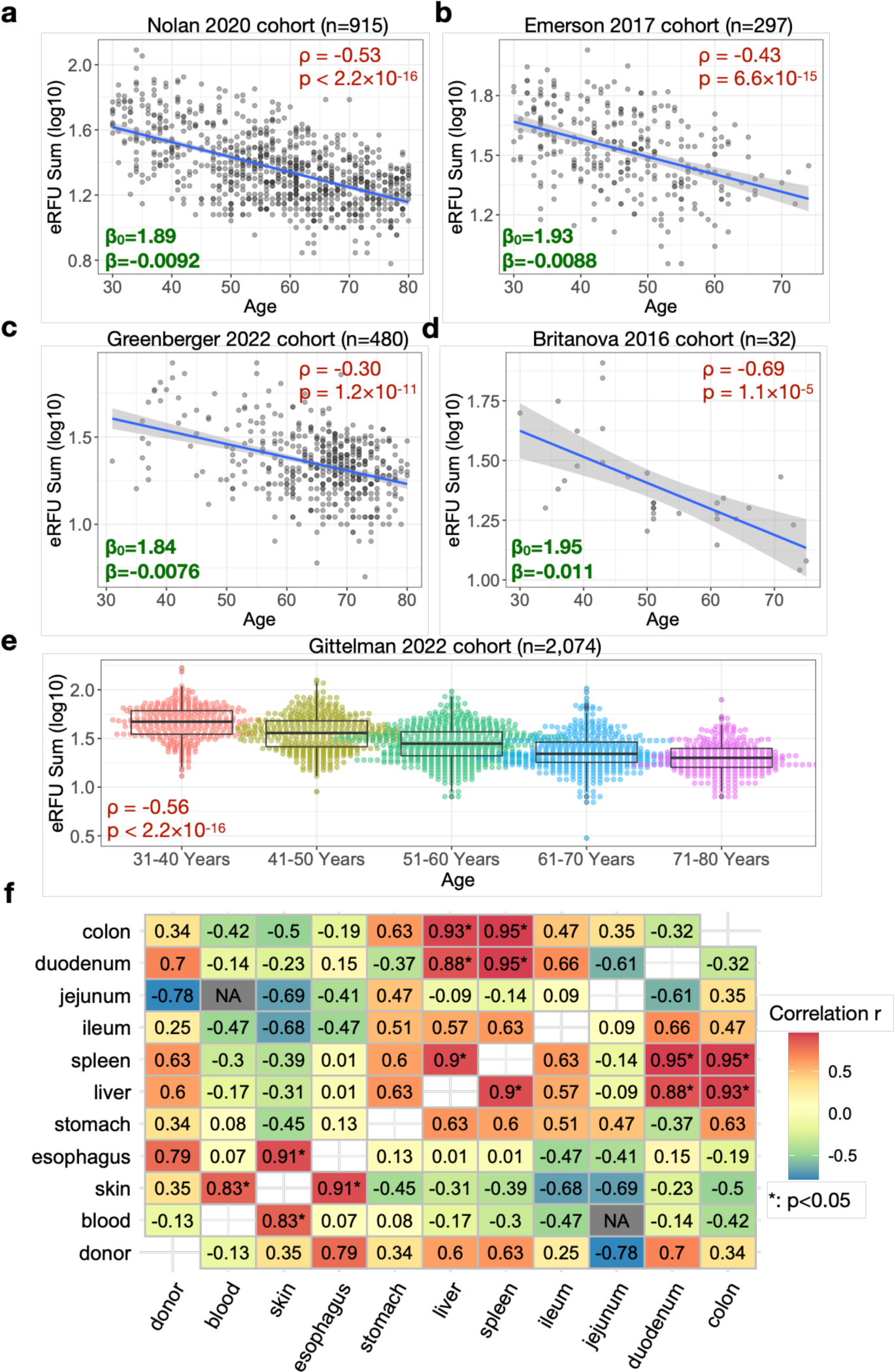
Quantification of eRFU loss with age in adult cohorts. **a**-**d**) Regression analysis of eRFU sum on age in four different adult cohorts. All individuals are within the range of 30 to 80 years. 95% confidence intervals of regression lines were calculated using 1.96 standard deviation estimated from the model. Top right text boxes show the Spearman’s correlation and p values between eRFU sum and age. **e**) Association of eRFU sum with age group information. Spearman’s correlation test was applied to evaluate statistical significance. **f**) Heatmap showing correlation of eRFU sums between a pair of tissue samples collected from 10 patients (postmortem). Statistical significance was evaluated using Pearson’s correlation test.

All above cohorts were derived from blood samples. We further investigated the aging trends of diverse organs using a postmortem patient cohort with TCR repertoires sequenced for skin, liver, spleen, and gastrointestinal (GI) track^44^. We observed that blood, skin and esophagus shared similar eRFU levels, whereas colon, liver and spleen were closely correlated (**Figure 2f**), though no p value passed FDR level at 0.05 due to small sample size.

### Accelerated loss of eRFUs under immunosuppressive conditions

We next investigated eRFUs loss among individuals with HCMV, HIV or COVID-19, in contrast to the healthy donors. For HCMV infection, HCMV+ was defined as patients with a positive serology. Interestingly, the age-association curves of patients with HCMV or SARS-CoV2 infections were similar to the healthy controls, while HIV infection was associated with a 37% drop in the overall eRFU levels (**Figure 3a**). Of the 13 eRFUs, 2428 and 4712 showed highest age-adjusted reduction in the HIV+ group (**Figure 3b**). These HIV patients (Towlerton 2020) received anti-retroviral treatment (ART)^45^, allowing us to investigate if therapeutic intervention could reverse this trend. Indeed, we observed a significant increase of eRFUs 2 years post ART (**Figure 3c**), which became more significant after correcting for participant age and individual baseline levels using a mixed effect model (**Figure 3d**). Based on these results, we speculated that immunosuppressive conditions lead to accelerated immune aging, manifested by faster reduction of the eRFU levels that could be rescued by therapy.

**Figure 3.**
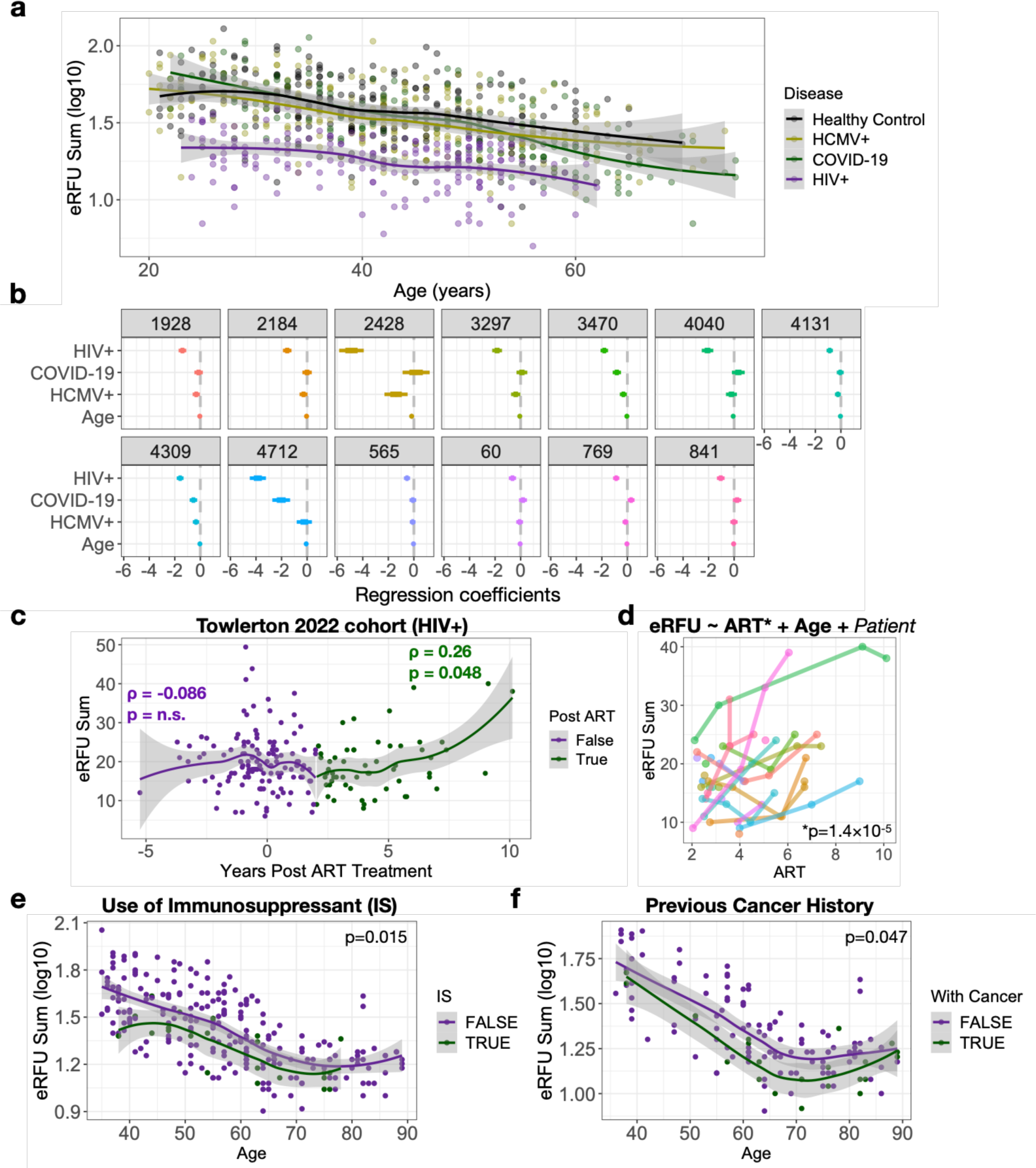
eRFUs are decreased in patients under immunosuppressive conditions. **a**) Scatter plot showing eRFU loss with age in different patient groups. **b**) Coefficient plot for all 13 eRFUs from linear regression with age and viral infection categories as covariates. **c**) eRFU sum dynamics pre- or post-ART treatment. Color change marks 2-year post ART. Spearman’s correlation test was implemented to estimate statistical significance. **d**) Dynamics of eRFU sum post ART treatment for each patient. Statistical significance was estimated using linear mixed effect model. **e-f**) eRFU dynamics with age stratified by the use of immunosuppressant (e) or existence of past cancer diagnosis (f). Statistical significance was estimated by logistic regression controlled for age.

To test this hypothesis, we leveraged the Nolan 2020 cohort, which comprehensively documented the medical records of over 1,400 COVID-19 patients. A subset of the patients reported usage of immunosuppressant drugs, such as corticosteroids to treat asthma^46^. We observed significantly reduced levels of eRFUs in those patients after adjustment for age (**Figure 3e**). In addition, 24 patients in this cohort were diagnosed with cancer before the time of blood draw. These patients may have been immunocompromised due to cancer-induced immune exhaustion^47^ or exposure to immunosuppressive therapies that are common for cancers^48^. Indeed, we observed a significant decrease of overall eRFU counts in these patients (**Figure 3f**). Interestingly, we noticed that use of immunosuppressant had a stronger influence among younger (35-50yrs) individuals, while cancer occurrence in the older (65-80yrs) patients was associated with higher reduction of eRFUs (**Figure 3e-f**). Together, our analysis indicated that immunosuppressive conditions might further lower the eRFU levels.

### Clinical impact of eRFUs in COVID-19 patients

To further understand the functional impact of eRFUs in the context of diseases, we performed a systematic analysis of TCR repertoire samples from several recent COVID-19 patient cohorts^22,42,49^. First, among the adult COVID-19 patients, higher overall eRFU level was associated with reduced risk of ICU admission when controlled for patient age (**Figure 4a**). Within hospitalized patients, we observed a suggestively higher eRFU count associated with fewer days in the hospital (**Figure 4b**). Quantitatively, at the same age, an increment of one eRFU clone reduced the odds of ICU admission by 6% and hospitalization by 0.29 days. Next, we investigated a cohort of pediatric patients^49^ who developed multisystem inflammatory syndrome in children (MIS-C), a rare but serious condition associated with COVID-19. Consistent with the analysis in adults, higher eRFU level in children is associated with lower risk of MIS occurrence (**Figure 4c**). Further, within the patients who developed MIS-C, higher eRFU count appeared to be associated with lower risk of cardiac involvement (**Figure 4d**), although this observation did not hold for neurologic involvement (**Figure S6a**). Notably, eRFUs have different correlations with age in adults (inverse) and children (positively), yet for both groups, eRFUs were associated with better clinical outcomes. Finally, we studied the outcome of COVID-19 mRNA vaccination in a patient cohort with hematologic malignancies^42^. Total eRFU count was positively correlated with anti-Spike protein antibody titer (**Figure 4e**; Spearman’s ρ=0.21, p=2.3×10^−6^), for both types of mRNA vaccines studied (BNT and Moderna) (**Figure S6b**). We controlled for related factors (age, gap between 1^st^ and 2^nd^ vaccines and COVID-19 status) using linear regression and confirmed that eRFU levels independently predicted higher antibody production after vaccination (**Figure 4f**). Together, our results suggested that higher eRFU counts were related to better outcomes in patients with COVID-19 and improved vaccine response.

**Figure 4.**
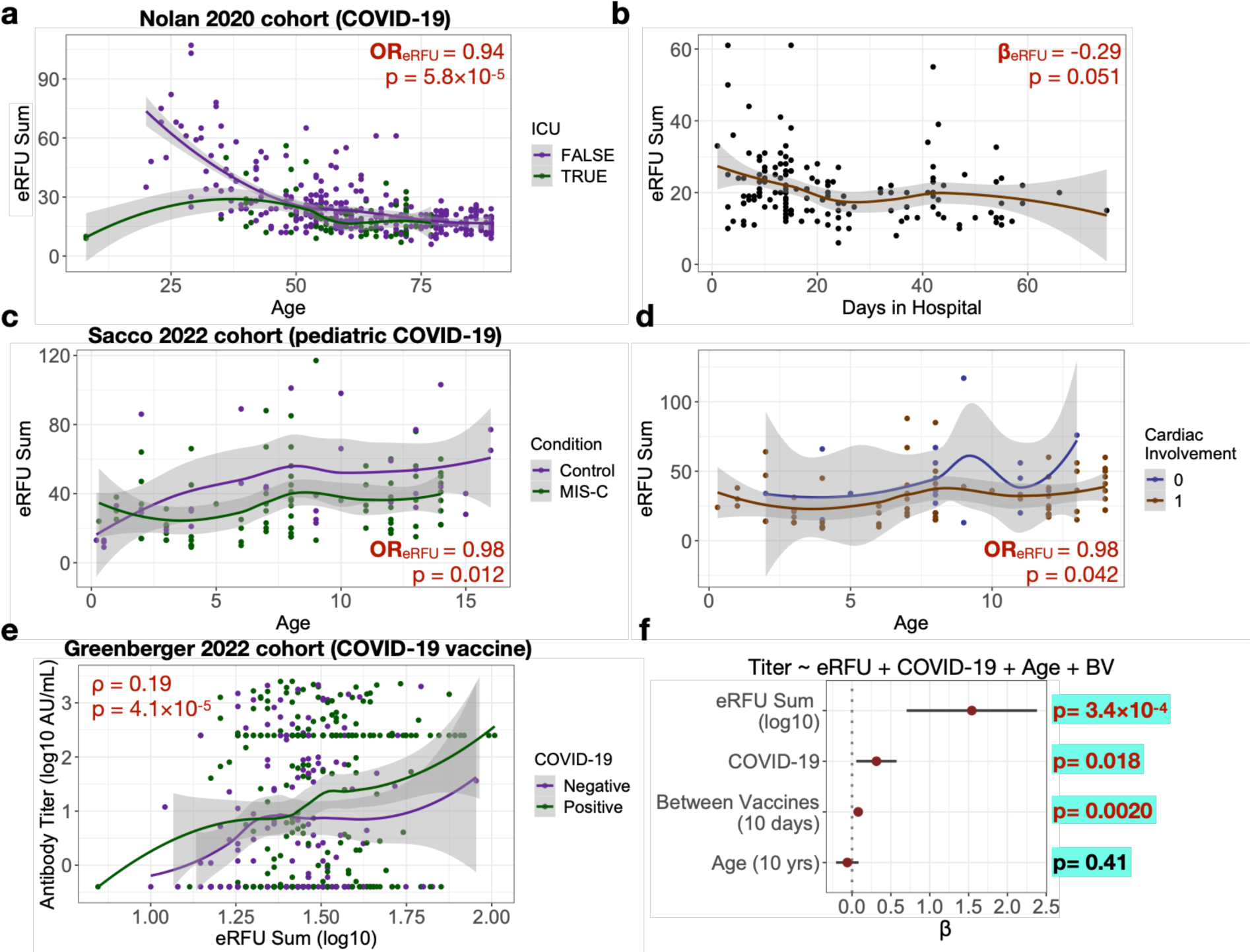
Clinical impacts of eRFUs in COVID-19 patients. **a**) Trending of eRFU sum with age stratified by ICU admission status. Odds ratio and p value were estimated using logistic regression controlled for patient age. **b**) eRFU sum trending with days in hospital. Linear regression controlled for age was used to estimate the impact of eRFU sum and p value. **c**-**d**) eRFU sum over age among pediatric COVID-19 patients stratified by disease groups: healthy control vs MIS-C (c) or cardiac involvement status (d). **e**) Scatter plot showing the relationship between eRFU sum and antibody titer after COVID-19 vaccination in an adult cohort. Statistical significance was evaluated using Spearman’s correlation test for both groups combined. **f**) Coefficient plot of linear regression analysis with related clinical covariates in the COVID-19 vaccination cohort.

### Secondary expansion of eRFUs following bone marrow transplantation

Given the potential predictive value of eRFUs in the clinical outcomes during acute infection, we next sought to investigate which factors influenced the expansion of eRFUs. For this purpose, we obtained TCR-seq samples from leukemia patients who received bone marrow transplantation, which allowed us to separately investigate the age associations of both donor and recipient. We first analyzed the Kanakry 2016 cohort^50^, including blood samples from 16 healthy donors and 46 recipients. We confirmed that recipient blood eRFU levels were negatively associated with age in the donors, and yet uncorrelated with recipient patient age (**Figure 5a**). In addition, using another cohort^51^ (Pagliuca 2021), we observed that in the patients who received peripheral blood stem cell (PSC) transfer, the association with donor age was no longer significant, in contrast to those who received bone marrow transplant (BM) (**Figure 5b**). These findings, together with our previous observations (**Figure S3-4**), suggested that senescence of the hematopoietic stem cells (HSC)^52^ may be a cause for the decreased eRFU levels in older populations with hematologic malignancies.

**Figure 5.**
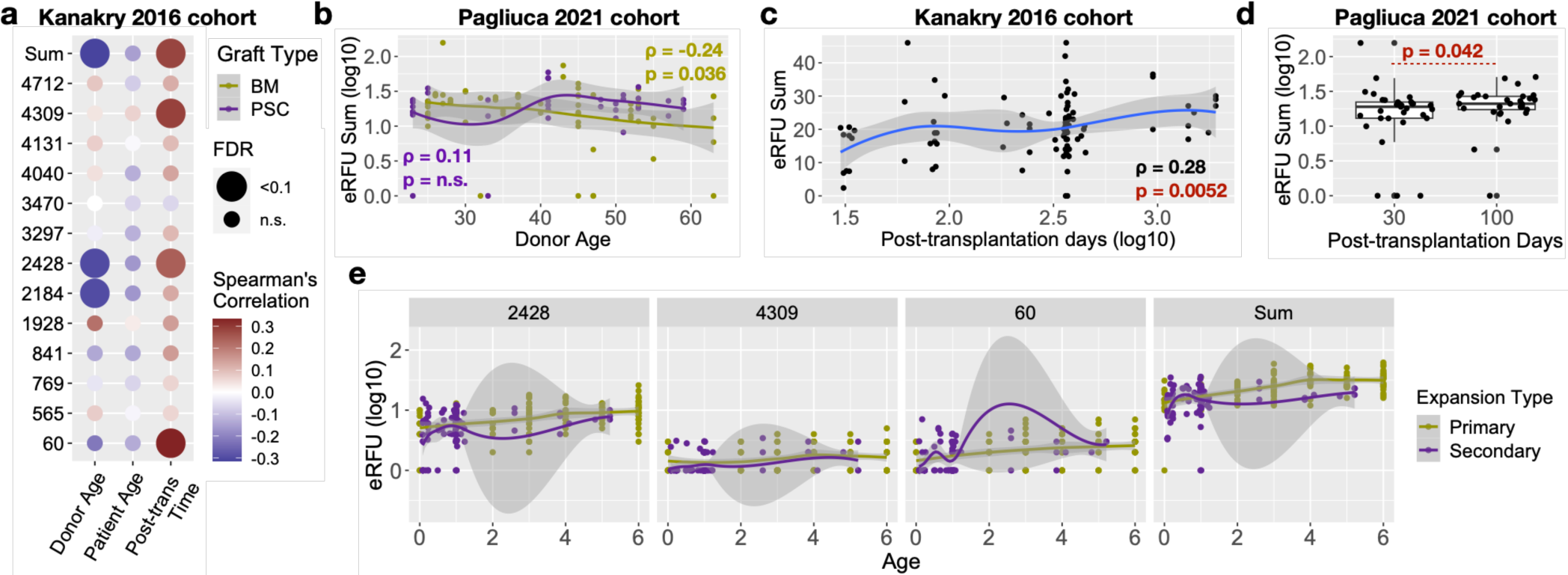
Secondary expansion of eRFUs after bone marrow transplantation. **a**) Heatmap showing the correlation of eRFU values and age (donors and recipient patients) or post-transplantation time (patients only). Statistical significance was evaluated using Spearman’s correlation test with FDR corrected using the Benjamini-Hochberg approach. Graft types included bone marrow (BM) or peripheral stem cell (PSC). **b**) Association of eRFU sum with donor age stratified by graft types. **c**) Increase of eRFU sum with post-transplantation days. For both (c) and (d), statistical significance was evaluated using Spearman’s correlation test. **d**) Boxplot showing eRFU increase after transplantation in Pagiluca 2021 cohort. One-sided Wilcoxon rank sum test was implemented to estimate statistical significance. e) Comparisons of primary and secondary expansions of selected eRFU clones or the summed value. Age (x-axis) was measured in years.

We next studied the changes of eRFUs after bone marrow transplantation. Interestingly, in both cohorts, we observed a significant increase of eRFUs by 2-fold within the first 100 days (**Figure 5c-d**). RFUs 60, 2428 and 4712 showed highest positive correlations with post-transplantation time (**Figure S7**). Notably, of these, RFUs 2428 and 4712 also displayed largest decrease in the Towlerton cohort of HIV patients (**Figure 3b**). We used the term ‘primary expansion’ to refer the increase of eRFUs after birth, and ‘secondary expansion’ the increase after bone marrow transplantation. We observed that even though selected eRFUs may reach a high magnitude in a secondary expansion, the overall count of eRFUs is lower than the primary expansion (**Figure 5e**). On average, the sum of eRFUs 5 years after transplantation was 59.6% of that in 5yr old children.

## Discussion

In this study, we identified quantifiable TCR repertoire units, or eRFUs, that robustly correlate with chronologic age, with a bell-shaped association peaking around 27 years old. Systematic investigation over 6,500 TCR-seq samples led to several intriguing discoveries: 1) immunosuppressive conditions, such as HIV infection or cancer, were associated with more loss of eRFUs than aging alone, and this trend is partially reversible with certain therapeutic interventions; 2) higher intrinsic eRFU levels were predictive of better outcomes during an acute viral infection in both pediatric and adult patients; 3) HSC transplant from younger donors might lead to a partial restoration of eRFUs in older recipients.

eRFU clones phenotypically belong to MAIT cells that recognize MR1-bound small molecules derived from bacteria^30^, which also are activated during viral infections^53,54^. Although the overall percentage of MAIT cells among CD3+ T cells in the blood is reported to dynamically change with age^55^, there has never been a systematic investigation of which MAIT clones have strong age associations. Our results suggested that only a subset of CD8+ MAIT clones expand after birth, and contract with aging. The age-associations of the 13 eRFUs defined using the Nolan 2020 and Emerson 2017 cohorts are repeatedly seen across all the TCR sample cohorts in our analysis. This is a surprising result, given the high diversity, high plasticity and potentially unseen batch effects of the immune repertoire samples. This reproducibility suggests that there might exist a driving immunologic factor, which could be the exposure to a common pathogen, a stimulating cytokine^56^ or an immune-related hormone^57^.

A likely explanation to the non-linear age-dynamics of eRFUs is that these T cell clones are expanded with antigen exposure during childhood, while contraction follows thymic involution^58^ and HSCs senescence^59^ in adulthood. A critical remaining question is, what antigens do these TCRs recognize? One hypothesis is that these RFUs are against common viral or bacterial infections given their constant exposures after birth. Through analyzing sequence logo and protein structure data, we identified 5-OP-RU as one potential bacterial target for eRFU 841. This finding is supportive for the above hypothesis. However, it should be noted that the pool of known antigen targets remains extremely limited, and it is unclear whether eRFUs, which mostly present in the form of CD8+ T cells, will directly kill the target cells to exert their functions.

Within the same age group, the total eRFU counts exhibited nearly 4-fold difference in the healthy donor cohort (**Figure 2e**). Genetics might be a contributing factor to this large variation, such as the polymorphism of the TRB variable gene region or the MHC loci^60^. Environmental factors, such as pathogen exposures, use of medications or supplements, lifestyle, disease history, psychosocial stress or other factors, could also be influential to eRFUs, though a systematic investigation is beyond the scope of this study. Notably, many of these factors, particularly diet, distress (e.g., depression), and smoking, among others, have been associated with other age-related phenotypes^61–64^. Leveraging epidemiologic studies will be critical in identifying these relationships.

There are several limitations of this study. First, eRFUs were identified using cross-sectional data. We do not know if within person changes over long age spans will replicate the patterns observed. Large longitudinal cohorts with blood specimens collected from multiple timepoints within the same individuals would be required to further evaluate the age-associations of eRFUs. Second, the eRFU analysis was focused on the βCDR3 region, which cannot fully determine the antigen specificity without the pairing of the ɑ chain. Future high-throughput single T cell sequencing efforts to pair with the ɑ chains is needed to understand the functional significance of the eRFUs. Finally, while multiple different cohorts were included in our analysis, their racial/ethnic diversity was limited. Also, many of the individuals in the analysis were experiencing diseases, which could influence the results.

In the future, more research is needed to investigate eRFUs and their potential clinical implications. The MR1-restricted eRFU clones are unlikely to be directly specific to SARS-Cov2 antigens, thus, their role in acute infections may transcend the type of pathogens involved, for example, boosting B cell responses through cytokine production^65^. Hence, it will be critical to understanding how the eRFUs are related to other metrics of immune system health and the potential role that they play in driving immunity in the context of infections and cancer. Additional validation in other populations as well as evaluating relationships with other age-related markers will be important to understand which aspects of the aging phenotype are captured by the eRFU measure. Finally, our findings might provide novel opportunities to develop approaches for immune rejuvenation, including vaccination approach to achieve higher eRFU levels^66^.

## Author contributions

B.L. conceived the project. B.L. and J.H. performed the analysis and wrote the manuscript. S.W. M.P and B.R. helped with data analysis.

## Acknowledgement

This work is supported by NCI R01 grants CA258524 (B.L.) and CA245318 (B.L.).

## Data and code availability

RFU codebase is available at: https://github.com/s175573/RFU. All the TCR repertoire sequencing data can be downloaded at https://clients.adaptivebiotech.com/immuneaccess with accession links listed in **Table S2**.

## Methods and Materials

### Datasets and preprocessing procedures

All TCR repertoire sequencing samples were accessed from the immuneAccess database (https://clients.adaptivebiotech.com/immuneaccess) managed by Adaptive Biotechnology by Nov 27^th^, 2023, or downloaded from literature-provided repositories. Samples collected from immuneAccess were profiled using the immunoSEQ platform developed by the company. Zip files were directly downloaded through the ‘Export’ function and selecting ‘v2’. Accession numbers for each cohort are available in Table S2. For each repertoire sample, sequences with missing variable genes or nonproductive CDR3 regions were removed. The top 10,000 TCRs with most abundant clonality were selected for RFU calculation. We exclusively used the top 10,000 clones to minimize the batch effects caused by differences in sequencing depths. These preprocessing criteria were applied to all the TCR-seq samples throughout this study. Associated clinical information for each cohort were individually accessed through literature. Single cell RNA sequencing paired with TCR sequencing data were downloaded from GEO by accession numbers GSE178991 and GSE194189.

### Repertoire Functional Units

The complete RFU method was previously described^67^. In brief, this approach divides into three major steps. First, we introduced a TCR embedding approach by clustering of over 20 million TCRs to obtain a trimer substitution matrix. Approximate isometric embeddings for each of the amino acid trimers were calculated based on multi-dimensional scaling. Embedding of each TCR sequence is calculated as the mean of all the embedding vectors from consecutive trimers. Next, in the trimer-embedding space, we defined TCR neighborhoods conservative across multiple individuals. This step is achieved by pooling 1.2 million TCRs from 120 healthy donors from a previous study^23^, and projecting them onto the Euclidean space with trimer-based embedding. We divided the TCR sequences in this space into 5,000 groups with the k-means method. We referred the centroid of each group as a ‘Repertoire Functional Unit’, or RFU. Last, to calculate the RFU vector of a new TCR repertoire sample, we select the top 10,000 most abundant TCRs based on clonal frequencies. We consistently used the top 10,000 clones for all the datasets to minimize the influence of sequencing depth. For each TCR, we calculate the embedding vector and assign it to the closest centroid from 5,000 RFUs. The value of each RFU is determined by the number of TCRs assigned to its centroid. We chose 5,000 as the group number so that the expected count for each RFU is 2.

### Regression Analysis

The relationship between eRFU sum (log10 scale) and age was quantified with ordinary linear regression for all the cohorts with numeric age information (Figure 2a-d). Constant variance of residuals (homoscedasticity) was visually confirmed on the residual plot using the Nolan 2020 cohort (Figure S3a). Linear regression with viral infection status and age was performed to evaluate the impact of each viral infection over eRFU sum (Figure 3b). Linear mixed effect model was implemented with patient age and time post ART therapy as fixed effect, and patient ID as random effect (Figure 3d). Regression was performed using R package *lme4* (v1.1-33). Statistical significance of ART over eRFU sum was evaluated using the *lmerTest* package (v3.1-3)^68^. Logistic regression controlled for patient age and/or other covariates was implemented to assess the impact of eRFU sum over binary clinical outcomes in COVID-19 patients (Figure 4). Odds ratio and statistical significance were both estimated from the R function *glm*.

### Public TCR analysis

4,425 public TCRs were defined as occurring in 90% of an age-representative population^12^. For each public TCR, we assigned its RFU number based on the method described above. To determine if eRFUs were enriched for the public TCRs, we randomly selected 13 RFUs (without replacement) from the 5,000 pool and calculated the number of public TCRs that had been assigned with an RFU within the random set. This number could be larger than 13 since it was the TCR count. The overlapping TCR count for eRFU was then compared to 1,000 random RFU sets to evaluate statistical significance. RFUs that were assigned with any public TCRs were compared to all or eRFUs for their age associations using the correlation values estimated in the Nolan cohort.

### HLA allele association analysis

We applied the Emerson 2017 cohort to investigate RFU association with HLA alleles (A or B loci) at 2-digits resolution. Specifically, we selected alleles occurring in 10 or more individuals in the full cohort. For each allele and for each RFU, we performed a one-sided t-test, to investigate if individuals with the allele have higher RFU count than those without, and recorded the z-score. We obtained and visualized the allele-by-RFU matrix of z-scores from this analysis (Figure S2a).

### Single cell RNA-seq data analysis for MAIT cells

An analysis of the gene expression pattern of MAIT cells was conducted using the Dong cohort^33^ (Access number: GSE178991) containing paired scRNA-seq and scTCR-seq data. We primarily used R/4.1.1 and R Package Seurat/4.0.5 for scRNA-seq data analysis. A total of 75,820 T cells passed quality control, with a mitochondrial gene percentage of less than 5% and a ribosomal gene ratio of less than 2%, allowing them to proceed to downstream analysis. When finding clusters, T cells were divided into 12 subgroups using a 0.3 resolution. T cells with age-associated RFU were highlighted in red in the tSNE plot and were found to be significantly enriched in cluster 7. The expression levels of four MAIT markers^34^ (TRAV1-2, CD161, ITGAE, and SLC4A10) were evaluated across all clusters. Notably, T cells in cluster 7 displayed typical MAIT gene signatures and were thus designated as MAIT cells.

For the Garner cohort^34^ (GSE194189), we directly downloaded the processed data from GEO which had been quality-controlled and normalized. This dataset contains paired scRNA-seq and scTCR-seq samples from flow-sorted MAIT cells and conventional memory T cells (Tmem). There were 12 donors in this dataset, aged from 27 to 65, comprising 89,456 cells in total. We further kept cells with only one pair of productive TCRɑ and TCRβ chains for both cell types. We further discarded some potential contaminating MAIT cells from Tmem cell population, which expressed TRAV1-2 paired with TRAJ33, TRAJ12 or TRAJ20 and had a 12 amino acid CDR3ɑ region. Finally, we retained 30,058 MAIT cells and 25,927 Tmem cells. We assigned RFU number to each T cell based on the TCRβ chain and obtained 1,971 T cells expressing eRFUs. Enrichment of eRFUs in specific cell subsets/states was performed using Fisher’s exact test. Differential analysis was done using the *FindMarker* function in Seurat package. Pseudotime trajectory was constructed using Monocle2^39^, and the direction of pseudotime was determined by CytoTRACE ^40^.

### Statistical Analysis

Computational and statistical analyses in this work were performed using the R programming language v4.3.0. FDR control was using the Benjamini-Hochberg method. Sequence logos were generated using package *ggseqlogo* (v0.1), by performing multiple sequence alignment (*msa*, v1.32.0) using CDR3s with length 16. Neighbor joining trees were calculated based on pairwise correlation matrix by RFU values and visualized using R package *ape* (v5.7-1). Subpanels of main figures were produced using *ggplot2* (v3.4.2). Loess smooth lines were estimated using the *loess* function in R with default parameters, which was automatically implemented in the *ggplot* function. Beeswarm plots were generated using R package *beeswarm* (v0.4.0). Violin plots were produced using R package *vioplot* (v0.4.0). Due to limited sample size (n=10), Pearson’s correlation test was implemented to calculate the eRFU associations between each pair of somatic organs and statistical significance (Figure 2f). Visualization of the heatmap was performed using *ggplot*.

---

**Figure S1.**
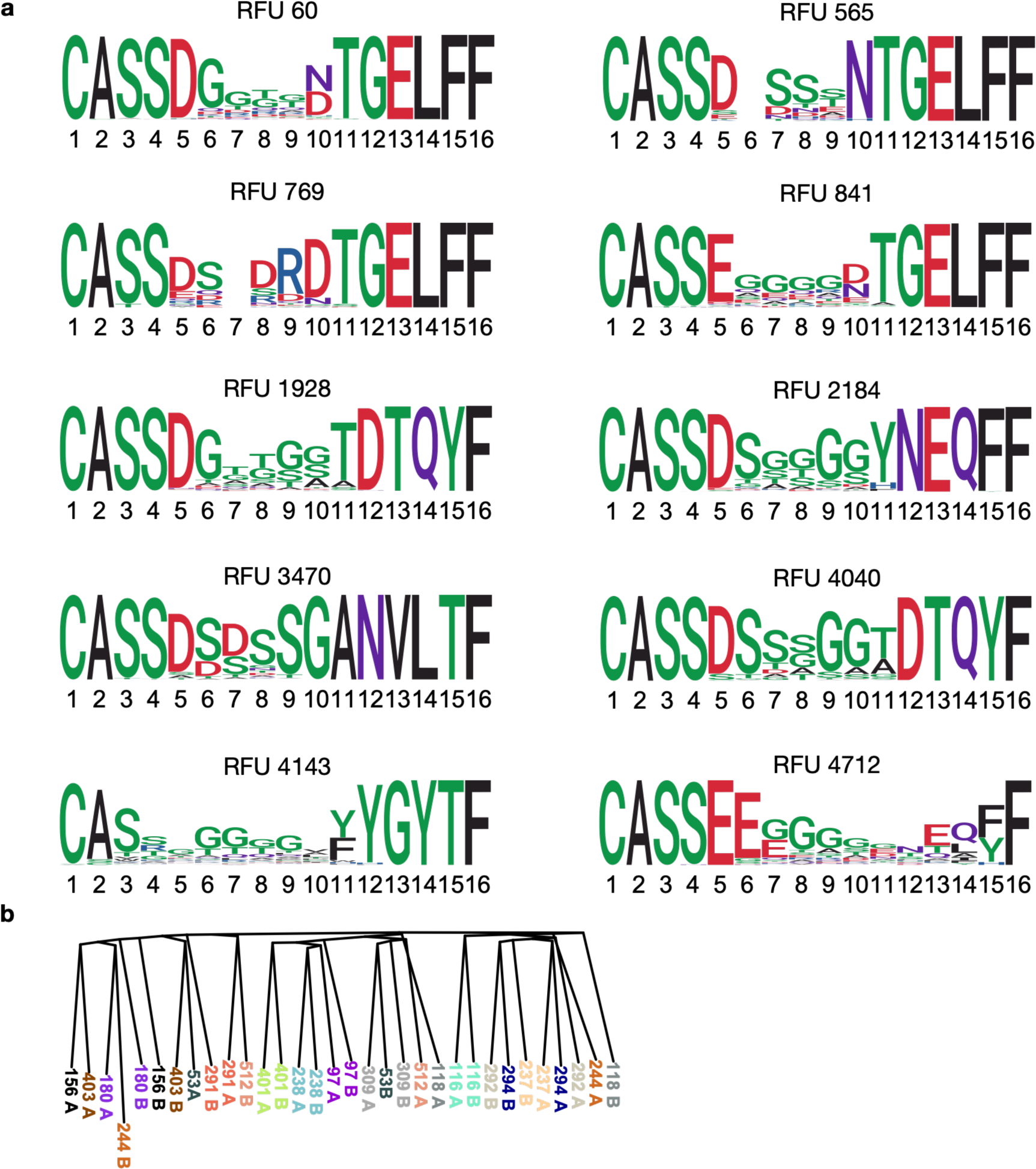
Additional features of eRFUs. **a**) Sequence logo plot for 10 eRFUs not shown in Figure 1. **b**) Neighbor joining tree showing the relationship between all the twins as in Figure 1h using all RFUs to calculate the distance matrix.

**Figure S2.**
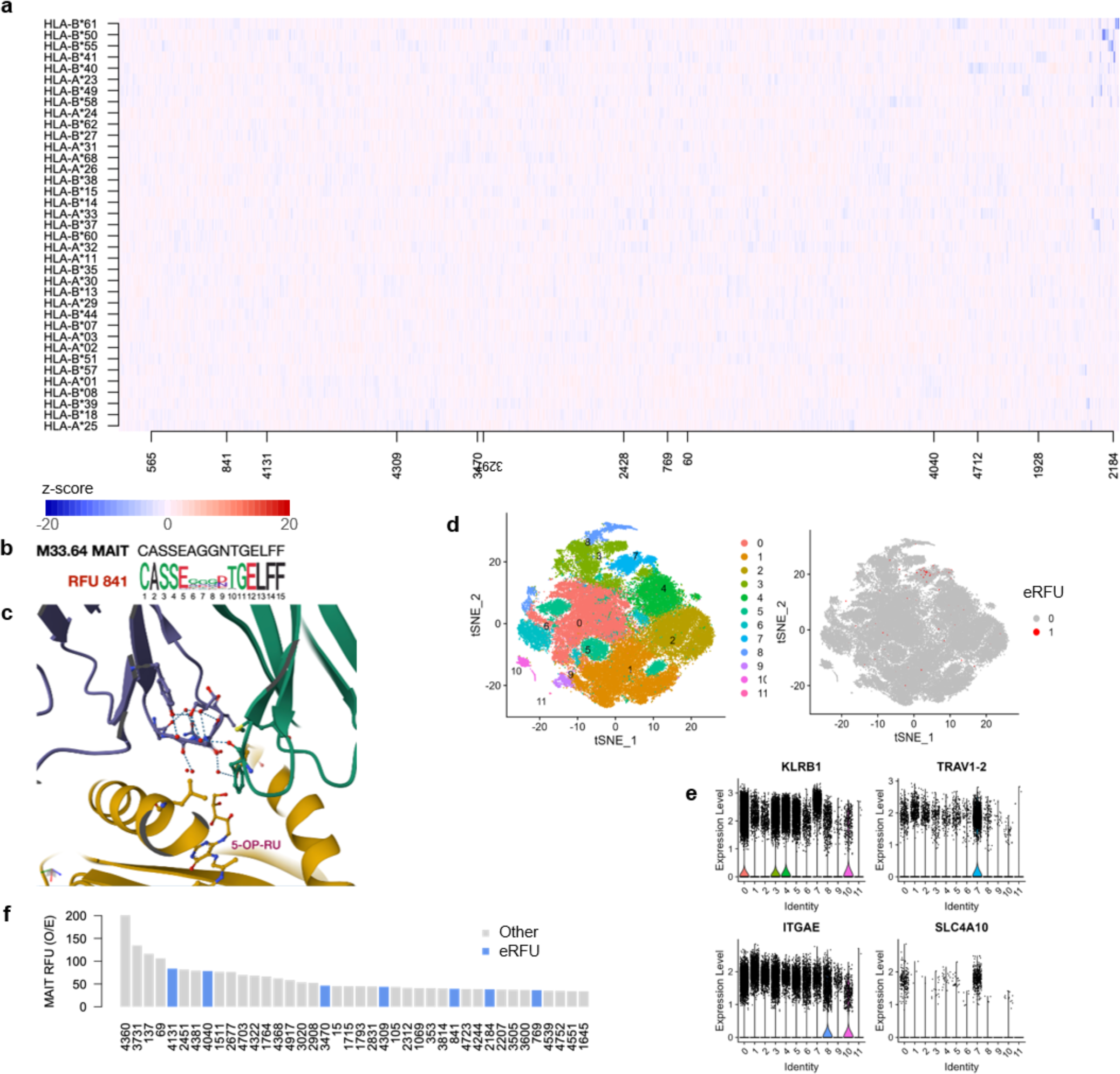
eRFUs displayed mucosal-associated invariant T cell signatures. **a**) Heatmap showing the z-scores of 5,000 RFUs and 37 HLA alleles. eRFU numbers are marked on x-axis. Details see Methods. **b**) Sequence motif comparison of the M33.64 TCR with RFU 841. **c**) Zoom-in view of the TCRβ chain (purple) of M33.64 interacting with MR1 (dijon yellow) and the antigen molecule (PDB accession: 5D7J). **d**) tSNE plot and clustering of T cells using single cell RNA-seq data (left) and same plot highlighting eRFU T cells with red color (right). **e**) Gene signature plot showing overexpression of selected MAIT cell markers in cluster 7. **f**) Enrichment analysis of MAIT RFUs among the T cells in the same single cell dataset.

**Figure S3.**
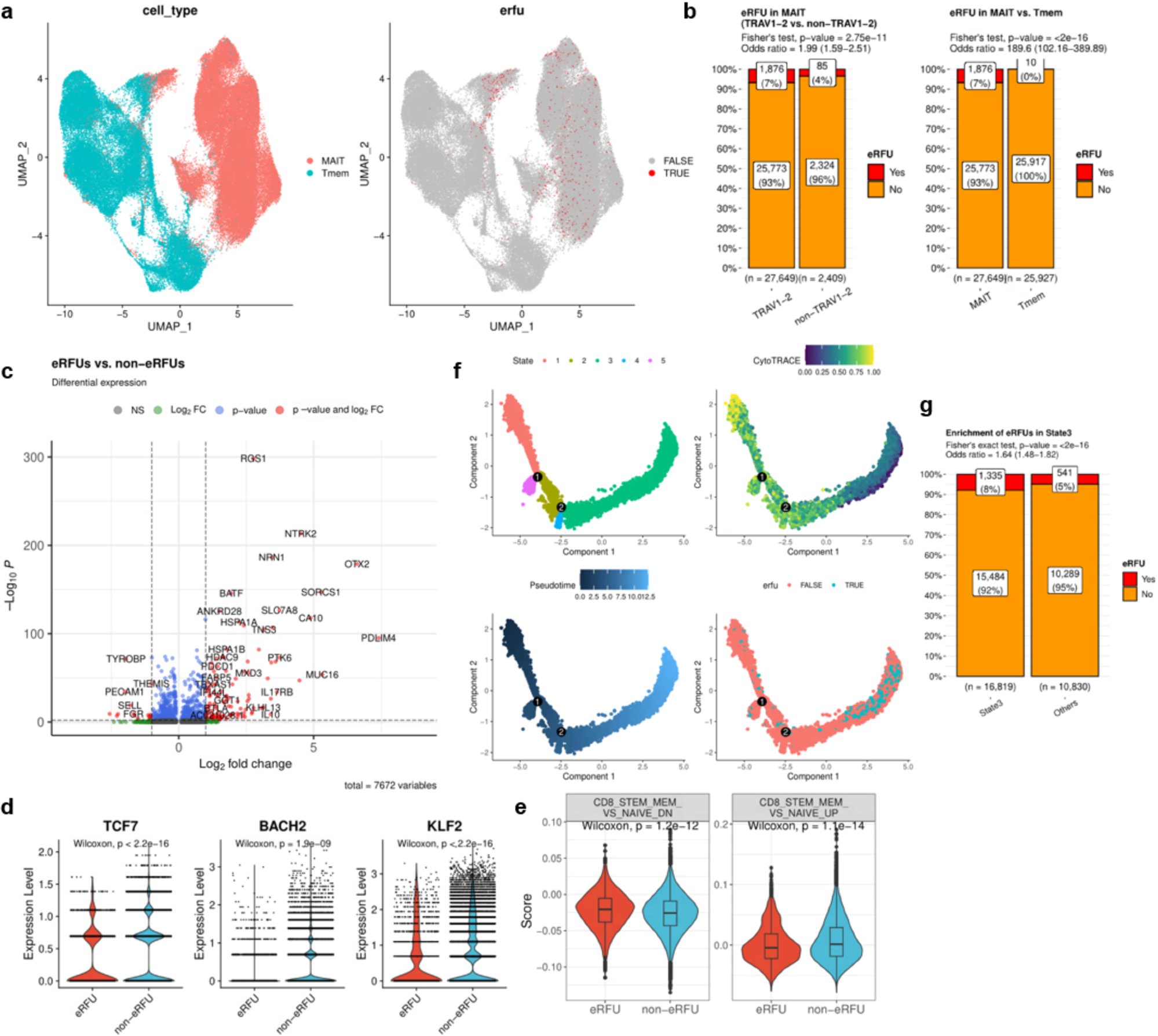
MAIT signatures and differentiation status of eRFU cells. **a**) UMAP plot showing the distribution of MAIT or memory T cells in the gene expression space (left) and the distribution of eRFU expressing T cells on the UMAP (right). **b**) Percentage plots for eRFU expressing T cells enrichment in TRAV1-2 vs non-TRAV1-2 (left) T cell subsets or in MAIT vs Tmem (right). **c**) Volcano plot showing the differentially expressed genes between eRFU and non-eRFU cells. Red color marks absolute log2 fold change greater than 1. Blue color indicates statistical significance at FDR=0.05. **d**) Violin plot showing the distributions of three putative T cell stemness markers. **e**) Violin plot showing the distribution of two scores measuring the signature genes that either down- (left) or up- (right) regulated in CD8 stem memory cells vs naïve CD8 T cells. **f**) Monocle pseudotime trajectory plot of all the MAIT cells with direction of differentiation determined by CytoTRACE, where lower value indicates higher differentiation status. **g**) Percentage plot showing eRFU cells enrichment in state 3.

**Figure S4.**
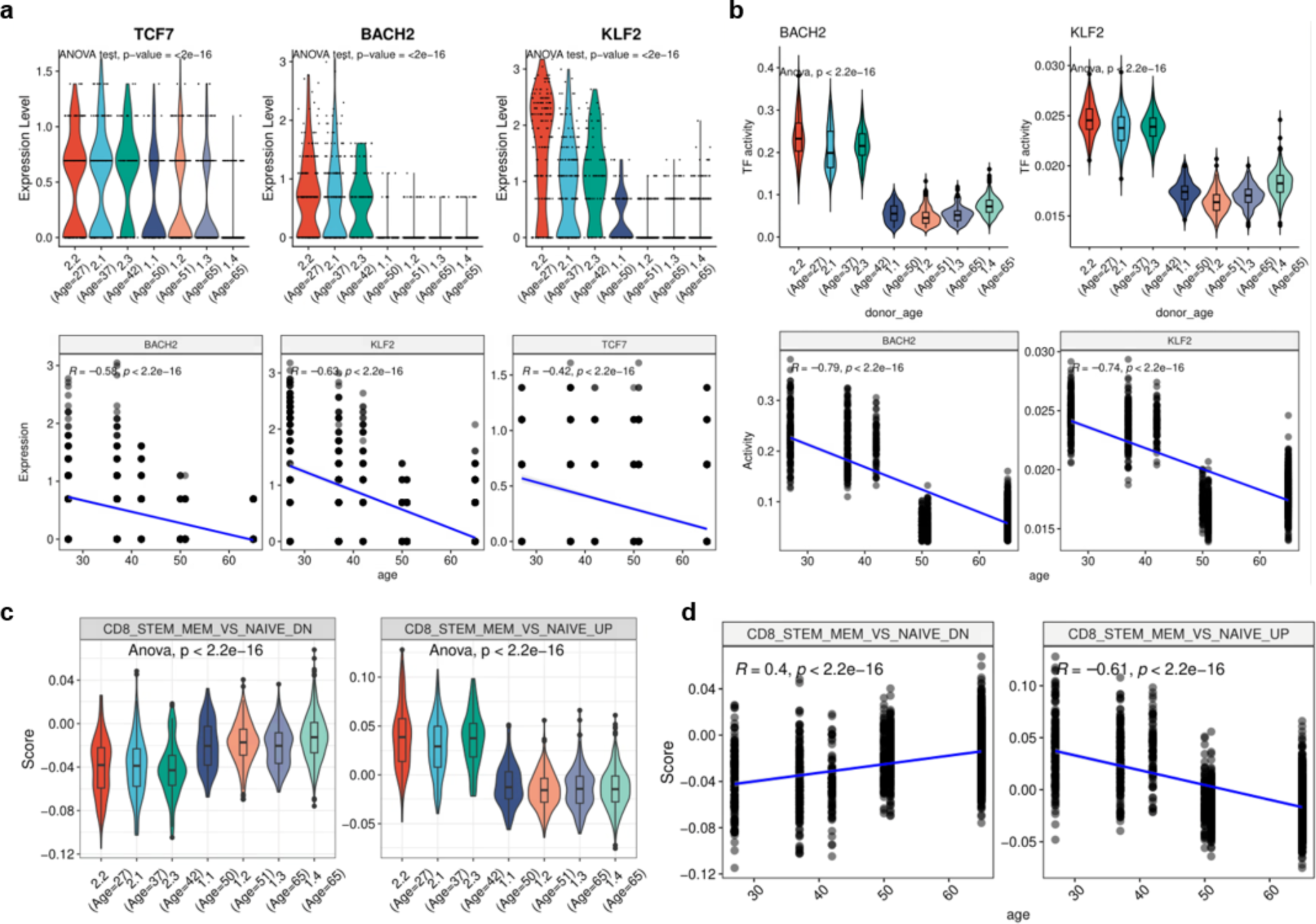
Age association of stemness markers in the eRFU cells. **a**) Violin (upper) or scatter (lower) plots showing the distributions of selected T cell stemness genes trending with age. Statistical significance in the upper panels was evaluated using one-way ANOVA, while the lower panels using Pearson’s correlation test. **b**) Similar analysis as in A) performed for transcription factor activity for stemness markers BACH2 and KLF2. **c-d**) Similar analysis performed for the scores measuring the signature genes down- or up-regulated in CD8 stem memory T cells compared to naïve T cells.

**Figure S5.**
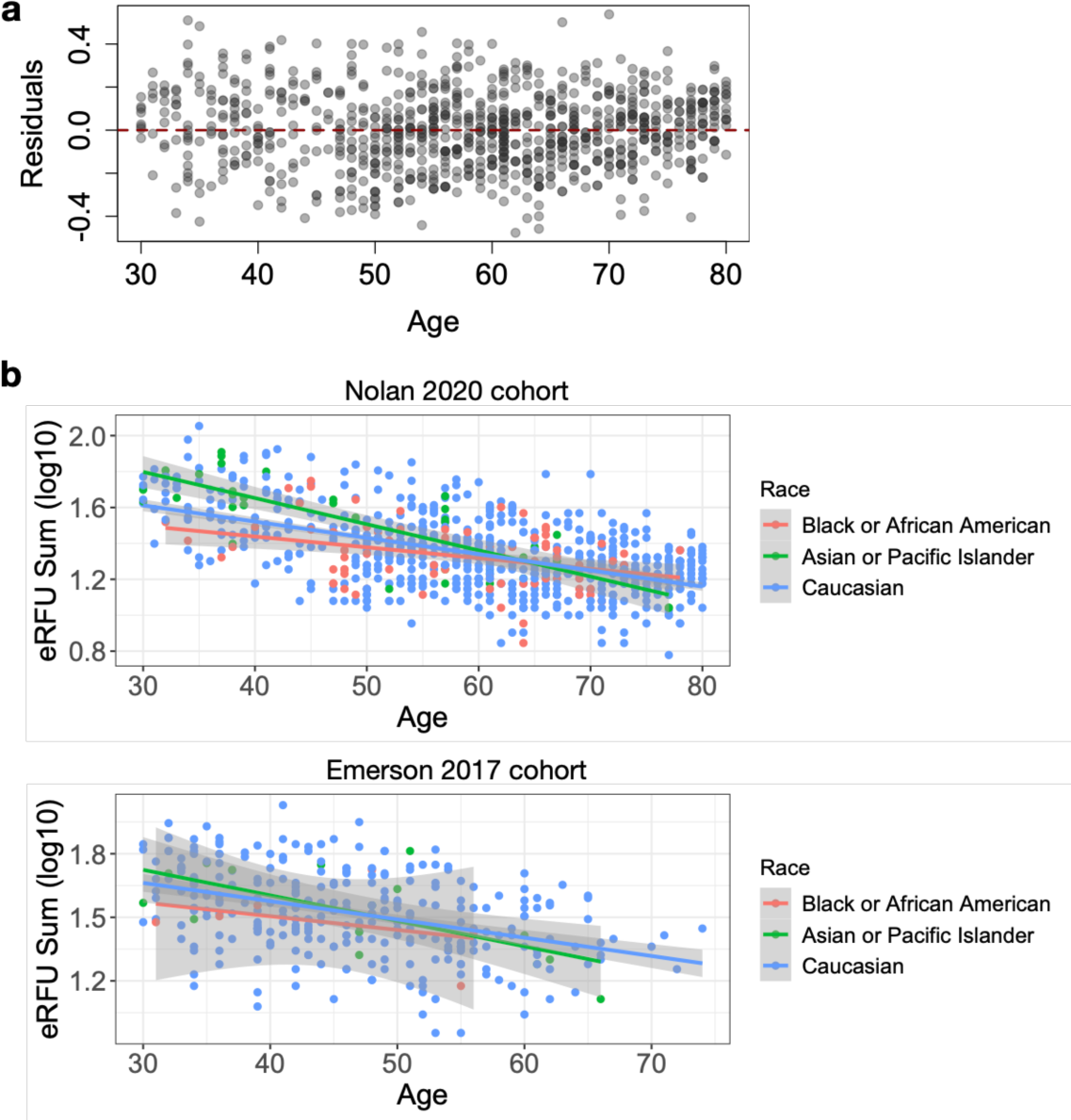
eRFU residual plot and distribution across different racial groups. **a**) Regression residual plot against age. Linear regression model log10(eRFU sum)∼Age was estimated using Nolan 2020 cohort. **b**) Trends of eRFU sum with age stratified by racial groups.

**Figure S6.**
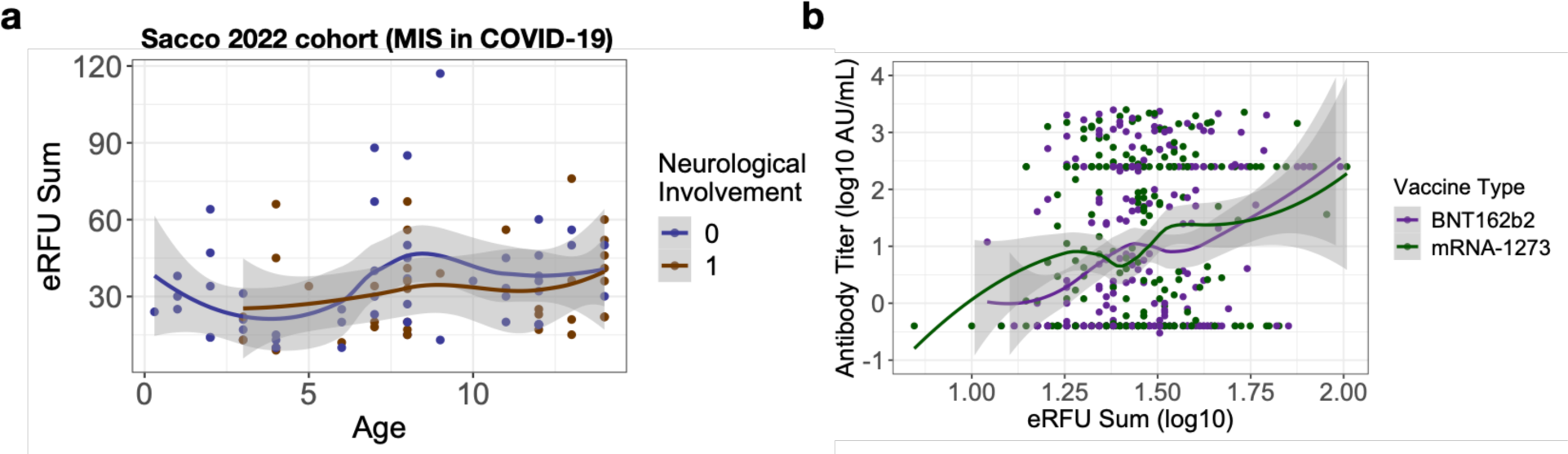
Additional results on eRFU functional impact. **a**) Scatter plot showing eRFU sum with age in pediatric COVID-19 cohort stratified with whether the patient developed neurological complications or not. **b**) Antibody titer increased with eRFU sum stratified by vaccine types.

**Figure S7.**
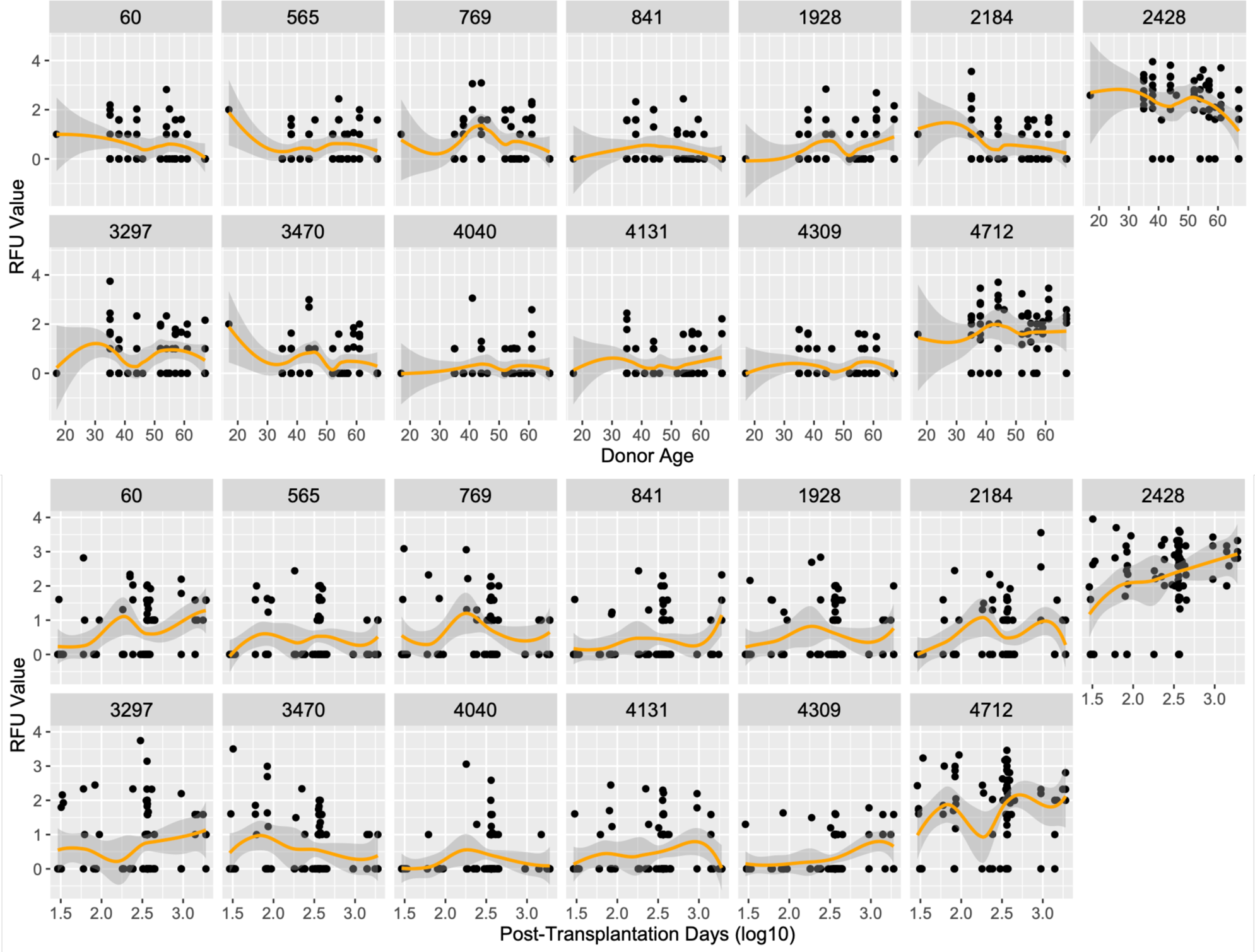
eRFU dynamics with donor age (upper panel) or with post-transplantation days (lower panel).

